# The ability of ddRAD-Seq to estimate genetic diversity and genetic introgression in endangered native livestock

**DOI:** 10.1101/454108

**Authors:** Ayumi Tezuka, Masaki Takasu, Teruaki Tozaki, Atsushi J. Nagano

## Abstract

Unplanned crossbreeding between a native livestock and a specific productive breed was one of the main reasons that caused the loss of valuable genetic resources in native livestock. To avoid further loss and damage of genetic resources in the native livestock, introgressed individuals should be distinguished to eliminate them by preventing any further employment in future mating plans. In general, the genetic diversity of native livestock had already decreased and mass elimination of introgressed individuals from the population endangers their existence. To solve this problem, high-resolution markers are required to discriminate between introgressed variation and native variation. Here, we applied ddRAD-Seq markers for native Japanese horse “Taishu” that has undergone recent genetic introgression. Genome-wide ddRAD-Seq markers can distinguish five breeds of native Japanese horses and Anglo-Arabian introgressed breeds. We found the signatures of genetic introgression of Anglo-Arabian at only two chromosomes; however, the signatures were separated in their genome suggesting that it might not be the cause of recent introgression. The genetic diversity of Taishu was less than other Japanese breeds and the decreasing genetic diversity is an urgent issue compared to genetic introgression. Although few signatures of recent introgression were detected, a lot of shared SNPs (10% of all SNPs in Taishu) were detected between Taishu and Anglo-Arabian. To avoid misestimation of the presence and degree of introgression in native livestock, information regarding shared SNPs and population genetic approaches need to be assessed by using the large number of genome-wide markers such as ddRAD-Seq.

## Introduction

The diversity and populations of native livestock breeds have dramatically decreased due to the dominance of a few breeds that have been selected for greater productivity (Rischkowsky and Pilling, 2007; Scherf and Pilling, 2015). Rapid and unplanned crossbreedings between native and productive breeds have frequently occurred, further depleting the valuable genetic resources of native livestock (Hanotte et al. 2010). To avoid further loss and damage to genetic resources of native livestock, introgressed individuals should be identified and removed. However, most endangered native livestock populations already have depleted genetic diversity, and the easy removal of introgressed individuals endangers their existence. Therefore, we should maintain a careful balance between preserving genetic diversity and removing genetic introgression by focusing at the genomic level (i.e., chromosomes and loci) to identify original genetic variations in native breeds and introgressed variations from other breeds. This could then be used to maintain the genetic diversity of native variants and eliminate the introgressed variants.

Genetic introgression is the movement of genes from one species into the genome of another via crossing. Therefore, genetic introgression can be difficult to detect without molecular data (Fitzpatrick et al. 2010; Ryan et al. 2009). The development of the Sanger-PCR methods has enabled estimates of genetic introgression using partial DNA sequences, such as mtDNA sequences and single sequence repeat (SSR) markers. However, detecting genetic introgression using partial DNA has some problems. For example, mtDNA is maternally inherited and does not recombine, so it can only be used to detect genetic introgression through females and cannot reflect the degree of genetic introgression. Meanwhile, SSR markers are usually no more than several hundred loci and incur the possibility of overestimating or underestimating genetic introgression. As genetic introgression has progressed for several generations, a larger number of markers is needed to accurately reflect the degree of introgression in individuals. Furthermore, individuals can be identified as hybrids from only a few markers even if it there is little introgressed variation in other regions. This can have a fatal effect on the conservation of native breeds because the few-introgressed individuals who carry native variations could be eliminated from the population, causing further decline in genetic diversity.

Endangered native livestock often have two problems: reduced genetic diversity and genetic introgression from other breeds. Removing genetic variation that has been introgressed from other breeds simultaneously removes the genetic variation of the native breed, further depleting the genetic diversity of native livestock breeds. Therefore, genome-wide, high-resolution markers are needed to achieve a suitable balance between removing the introgressed genetic variations and retaining the genetic diversity of native livestock.

Double-digested restriction site-associated DNA sequencing (ddRAD-Seq) is used to obtain thousands of single nucleotide polymorphisms (SNPs) across the genome (Baird et al. 2008; Peterson et al. 2012). To detect genome-wide SNPs, various molecular methods have been used to generate libraries for use in next-generation sequencing (Futschik and Schlotterer, 2010; Mardis, 2008; Nielsen et al. 2011). One effective method is ddRAD-Seq, a variation of genotyping-by-sequencing (GBS) (Poland and Rife, 2012). In this strategy, genomic DNA is fragmented using restriction enzymes and sequenced using next-generation sequencing technologies to obtain SNPs that are located next to target restriction sites. As there are far fewer sequences, this strategy increases the coverage of fragments and provides reliable data for many samples. There are some favorable aspects for applying ddRAD-seq to endangered native livestock. For example, ddRAD-seq can be applied to all target species and is cost-effective compared to whole-genome sequencing and SNP chips. Although SNP chips designed for some livestock can provide tens of thousands of accurate SNPs, it is possible that the SNPs do not include those of native livestock. Also, SNP chips are not readily available for some native livestock, whereas ddRAD-seq can be applied to almost all native livestock without additional experiments. Thus, it is clear that ddRAD-Seq is useful for detecting the genetic introgression of native livestock and is a versatile method for genetic introgression problems for many native livestock.

ddRAD-Seq has often been used to detect genetic introgression and hybridization in wild species (Chattopadhyay et al. 2016; Combosch and Vollmer, 2015). Most of the previous studies aimed at finding the geographical hybrid zone and detect signatures of hybridization and backcrossing between target species. In contrast, ddRAD-seq approaches for endangered native livestock aimed to identify introgressed individuals and decide whether the individuals were needed to be included in the conservation plan. As native livestock breeds have lower genetic diversity compared to wild species, introgressed loci and regions should be identified using high-resolution markers to create sustainable conservation plans. Here, we confirmed the ability of ddRAD-Seq to identify introgression variations with high resolution.

In this study, we examined native Japanese horse breeds on Tsushima-island in Nagasaki prefecture, Japan. The breed Taishu is listed by the FAO as “critical maintained” (Rischkowsky and Pilling, 2007; Scherf and Pilling, 2015). Taishu was introgressed with the Anglo-Arabian breed for military use during World war II (Hayashida, 1972). There is documented evidence of this genetic introgression, which describes one main event of crossing between these two breeds (Hayashida, 1972). This single introgression of Taishu presents a good opportunity to estimate genetic diversity and genetic introgression in native livestock using ddRAD-Seq. In addition, mtDNA sequences, SSR markers, and SNP chips for horse (*Equus*) are available. SNP chips for horse can produce 54 000 and 670 000 SNPs (McCue et al. 2012; Schaefer et al. 2017). Some studies used these methods in some native Japanese horses. Therefore, we can compare the results of our ddRAD-Seq with those from previous studies. We obtained about 10 000 SNPs by ddRAD-seq and identified genetic introgression and native loci on each chromosome in Taishu breed. We demonstrate the utility of ddRAD-Seq for the conservation of endangered native livestock breeds.

## Material and methods

### Sample collection and DNA extraction

We collected fresh blood samples from 57 individuals of 5 different breeds of Japanese native horse (*Equus caballus*), Taishu (N = 38), Kiso (N = 5), Miyako (N = 9), Yonaguni (N = 5), and Hokkaido (N = 6) in Japan. There are records that Taishu, Kiso, and Hokkaido were introgressed with European breeds, whereas Miyako and Yonaguni are not introgressed. Taishu is genetically closer to the Kiso-Hokkaido clade than to the Miyako-Yonaguni clade (Tozaki et al. 2003). In addition, blood samples from the Anglo-Arabian breed (N = 5) that was introgressed into Taishu, were provided by Goryo Bokujo (Imperial Stock Farm). None of these breeds have genetic crossing between them at present. The blood samples were collected in a tube with EDTA and kept at −20 °C until DNA extraction. Total genomic DNA was extracted from the whole blood using the Maxwell 16 Blood DNA Purification Kit (Promega, USA).

### DNA sequencing of mtDNA

16S rRNA sequences of mtDNA were amplified using primers designed by Achilli et al. (2012). All amplifications followed PCR protocols for a reaction volume of 10 μl: 100 ng of the DNA template, 5.0 μl 2: KAPA, and, 0.2 μM each primer. The amplification conditions were as follows: 2 min at 98 °C; then 30 cycles of 30 s at 98 °C, 30 s at 66 °C, and 1.5 min at 72 °C; ending with 15 min at 72 °C. The PCR products were purified and cleaned using ExoSAP-IT Express PCR Product Clean-UP (Affymetrix). Sequencing was performed by Macrogen (Seoul, Korea). Sequencing was performed using nested primers as designed by Achilli et al. 2012. The obtained sequences were aligned using Clustal X software (Larkin et al. 2007).

### Library preparation and sequencing in ddRAD-Seq

Library preparation was composed of 5 steps. First, restriction enzyme digestion and adapter ligation were performed in 10 µL reaction mix: 2 µL sample DNA (20 ng/µL), 0.5 µL EcoRI (10 U/µL, Takara, Osaka, Japan), 0.5 µL BglII (10 U/µL, Takara), 1 µL 10x NEB buffer 2 (New England Biolabs, Ipswich, MA, USA), 0.1 µL 100x BSA (Takara), 0.4 µL EcoRI adaptor (5 µM), 0.4 µL BglII adaptor (5 µM), 0.1 µL ATP (100 mM), 0.5 µL T4 DNA Ligase (600 U/µL, Enzymatics, Beverly, MA, USA) and 4.5 µL nuclease-free water. The digestion and ligation were performed at 37 °C for 16 h.

The two Y-shape adapters were prepared by annealing two partially complementary oligo-DNAs. A mixture of 100 µM adapter F and R was annealed using a thermal cycler with the following program: 95 °C for 2 min, slow-cooled to 25 °C (0.1 °C/s), followed by 30 min at 25 °C. The annealed adapter (50 µM) was stored at −20 °C. It was diluted to the working concentration (0.4 µM) just before use. The oligonucleotide sequences of the Y-shaped adaptors were as follows: BglII_adaptor_F: 5′ ACG GCG ACC ACC GAG ATC TAC ACT CTT TCC CTA CAC GAC GCT CTT* C*C-3′ BglII_adaptor_R: 5′-G*A*T CGG AAG AGC TGT GCA GA*C* T-3′; EcoRI_adaptor_F: 5′ A*A*TTGAGATCGGAAGAGCACACGTCTGAACTCCAGTC*A*C ‐3′, where *signifies a phosphorothioate bond and “/Phos/” signifies a phosphorylation.

The ligation product was purified using AMpure XP beads (Beckman Coulter, Brea, CA, USA) as follows: 10 µL of the AMpure XP and 10 µL ligation product were mixed by pipetting and kept at 25 °C. for 5 min. The purification was performed according to the manufacturer’s instructions. Then, the purified adaptor-ligated DNA was subsequently amplified by PCR. Amplification was performed in 10 µL reactions: 2 µL DNA, 2 µL Index primer (5 µM), 1 µL TruSeq_Univ_primer (10 µM), 5 µL 2X KAPA HiFi HS ReadyMix (KAPA Biosystems). The PCR was executed with 94 °C for 2 min and 20 cycles of 98 °C for 10 s, 65 °C for 15 s, and 68 °C for 15 s. After PCR, the product was preserved at 4 °C. The oligonucleotide sequences of the primers were as follows: TruSeq_Univ_primer: 5′; and Index primer: 5′ GAC GGC ATA CGA GAT XXX XXX GTG ACT GGA GTT CAG ACG TGT-3′ signifies an index sequence.

The PCR products of all samples were combined and concentrated using AMpure XP beads. The combined PCR product was mixed with equal volume of AMpure XP. Then, the mixture was placed on a magnet, and after 5 min its supernatant was removed. The remained beads were washed by adding 75% EtOH in excess volume than the mixture and removing the supernatant after 30 s; this was repeated twice on the magnet. After addition of 50 µL nuclease-free water, the beads were resuspended by pipetting and kept at 25 °C. for 1 min. The concentrated DNA was obtained by collecting the supernatant. The concentrated DNA was purified by size selection using E-Gel SizeSelect 2% agarose (Life Technologies, Carlsbad, CA, USA). Approximately 350 bp fragments were retrieved; their concentration was measured using a QuantiFluor dsDNA System (Promega, Madison, WI, USA), and the quality was measured with a Bioanalyzer DNA HS kit (Agilent Technologies, Santa Clara, CA, USA). After preparation of the library, 50-bp sequences of the *Bgl*II digested side of the DNA fragments were read using a HiSeq2000 and Hiseq2500 (Illumina, San Diego, CA, USA) by Macrogen. The sequenced reads were demultiplexed by CASAVA 1.8.2 (Illumina). Fastq files were deposited into the DNA Data Bank of Japan Sequence Read Archive as accession no. DRA007047.

### SNP calling

After removing the reads that contained low-quality bases and adapter sequences from the raw sequence reads using Trimmomatic ver. 0.33 (Bolger et al. 2014), SNPs were called with Stacks ver. 1.37 (Catchen et al. 2013). This process was performed with the default settings of the pipeline *ref_map.pl* in Stacks (population analysis, ‐m 3 ‐M 2 ‐n 1). We then generated an HTML report using a program *Stacks binder* (Yasugi et al. 2018) to visually check the summary of the ddRAD-Seq library and the results of SNP calling (Supplemental Material S1).

### Genome-wide locus-based phylogeny

We also reconstructed the phylogeny of the genome-wide ddRAD-Seq data set in RAxML (Stamatakis, 2014). We used the GAMMA+P-Invar model of sequence evolution and performed a single full maximum likelihood tree search. We applied that the rapid bootstrap algorithm with 1000 replicates to each data set. The resultant tree was plotted using Figtree (http://tree.bio.ed.ac.uk/software/figtree/).

### Mitochondrial DNA-based phylogeny

We aligned 1828 bp in lengths, including 16S rRNA sequences of 53 samples, and reconstructed the phylogeny of 16S rRNA sequences using maximum likelihood in RAxML, version 8.2.7 (Stamatakis, 2014). We used the GAMMA+P-Invar model of sequence evolution and performed a single full maximum likelihood tree search. We applied the rapid bootstrap algorithm with 1000 replicates to each dataset. The resultant tree was plotted using Figtree (http://tree.bio.ed.ac.uk/software/figtree/).

### Test of genetic introgression using allele frequency data

We analyzed 9 609 SNPs using TreeMix ver. 1.12 software (Pickrell and Pritchard, 2012), which were used to infer population history, including divergence and gene flow, using allele frequency data under genetic drift. TreeMix showed the maximum-likelihood tree with estimated hybridization events, including the direction of gene flow. We ran TreeMix with 0 and 1 migration events from Anglo-Arabian breeds to Taishu breed with various migration rates.

We used a model-based clustering approach in STRUCTURE (Pritchard et al. 2000) to identify genetic clusters within Japanese native horses and to investigate the degree of genetic introgression at individual level. We ran STRUCTURE from K = 1 to 6, with 10 iterations per K by using all individual data. Then, we ran STRUCTURE from K = 1 to 3, with 10 iterations per K by using data of Anglo-Arabian individuals and Taishu individuals. Each iteration included a burn-in of 50 000 generations, followed by MCMC for 100 000 generations. We obtained the optimal K using methods in Evanno et al. (2005) (EVANNO et al. 2005) the STRUCTURE harvester (Earl and vonHoldt, 2012). We plotted the results using STRUCTURE PLOT version 2.0 (Ramasamy et al. 2014).

To test if variations shared SNPs, the Taishu and Anglo-Arabian breeds could be explained by genetic introgression, rather than ancestral variation, we performed ABBA/BABA tests by calculating D-statistics and Z-scores. This measured the signatures of alternative phylogenetic asymmetry and the proportion of the genome that was shared between the two breeds due to genetic introgression. We defined 1 of the 4 Japanese native breeds and Taishu as sister clades. “Anglo-Arabian” and “Thoroughbred” were assumed as the introgression breed and the outgroup breed, respectively. For the test, we used the “CalcD” and “WinCalcD” functions in the “evobiR” R package (Blackmon et al. 2013), with 1000 replicates for each individual test and 100 replicates for each chromosome and regions to estimate variance.

### Linkage disequilibrium in Taishu populations

Before detecting regions of high linkage disequilibrium (LD) in the Taishu breed, we conducted phasing of genotype data and imputation of missing data using Beagle ver. 5 (B. L. Browning and S. R. Browning, 2016). We used the phased data with Haploview software ver 4.2 (Barrett, 2009) to calculate pairwise measures of LD among SNPs on the same chromosomes. Using the default method, we divided the region into blocks of strong LD using a standard block definition based on confidence intervals for strong LD (Gabriel et al. 2002) and minor allele frequencies > 0.05. If the haploblock size was > 10 kb, we defined the regions with variations shared between Taishu and Anglo-Arabian as potentially introgressed variation.

## Results

### SNPs detection in the native Japanese horse

We obtained 312 035 915 sequence reads after removing undesirable reads. The median read number per sample was 4 588 763 (interquartile range: 2 747 437–6 587 774). The median average quality value per sample was 37.465 (interquartile range: 36.77–37.63). RAD-seq library summary is reported in Supplemental Material S1. From these reads, Stacks was used to build 1 363 411 loci that contained 0 or more than 1 SNP. Supplemental Material S1a and A.1b show the number of loci shared by individuals. The number of loci decreased as the number of matching samples increased, with or without SNPs. All information regarding the reads used in Stacks, including that described above, is given in Supplemental Material S1a. We used 9 609 SNPs for analysis, filtered using the criteria over 75% matching samples, 1 SNP, 2 alleles, cut off 0.05% minor alleles, and mapped them to the chromosomes.

### Genetic introgression at the population level - Maximum Likelihood phylogenies of 5 native Japanese breeds and Anglo-Arabian -

To confirm admixture between Anglo-Arabian population and Taishu population, we reconstructed the ML phylogeny using 9 609 SNPs. The phylogeny showed that each breed consisted of a single clade (Fig. 1). All individuals of Taishu were grouped in a single clade and separated from all individuals of Anglo-Arabian. Some population structure was detected in the Taishu clade and the genetic distances between Taishu subpopulations were relatively large, but the genetic distances in both subpopulations of Taishu and Anglo-Arabian were large. We sequenced 1829 mtDNA sequences, including 16S rRNA that have slowest evolutionary rate of the present *Equidae* (Achilli et al. 2012). We detected 8 SNPs in 16S rRNA of mtDNA. The phylogeny constructed using mtDNA did not show the pattern of breeds (Supplemental Material S2.).

**Fig. 1.**
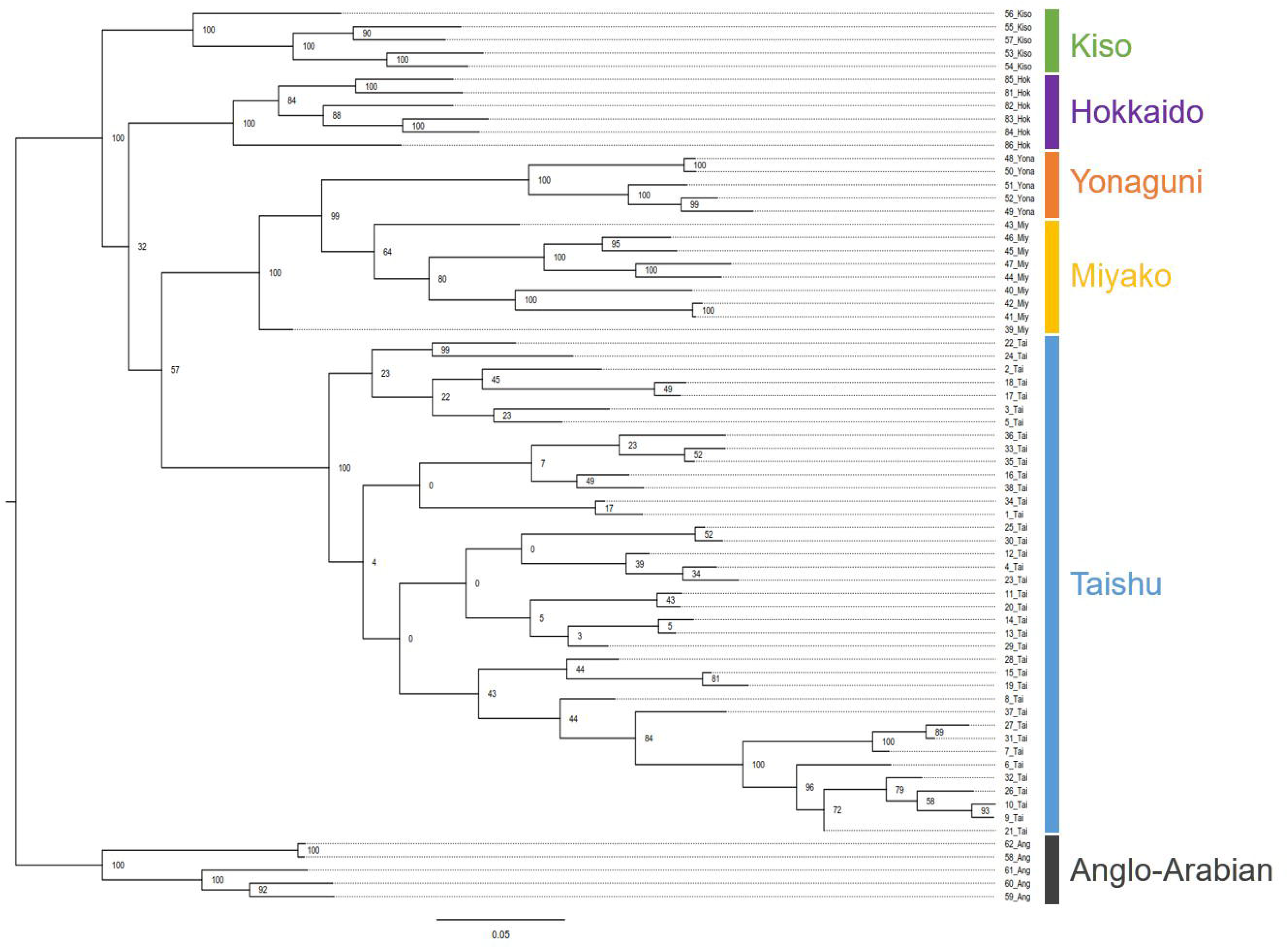
Phylogeny of 5 Japanese horse breeds and Anglo-Arabian. Maximum Likelihood phylogeny of genome-wide SNPs. The color boxes represent different breeds. Green, purple, yellow, orange, blue, and gray represent Kiso, Hokkaido, Yonaguni, Miyako, Taishu, and Anglo-Arabian, respectively.

### Genetic introgression at the population level - TreeMix analysis -

To estimate a migration rate of the genetic introgression from Anglo-Arabian into Taishu, the TreeMix approach was applied to all 6 breeds. A tree with 0 edges (i.e., no introgression event) explained 98.55% of the variance in relatedness between breeds (Fig. 2a). When a 4% migration edge was allowed from Anglo-Arabian into Taishu, the variance in relatedness between breeds was explained by the model that reached 98.80 %, which was the best-explained migration weight at an introgression from Anglo-Arabian to Taishu (Fig. 2b), based on the assumption that migration did not dramatically improve the variance in relatedness of the tree.

**Fig. 2.**
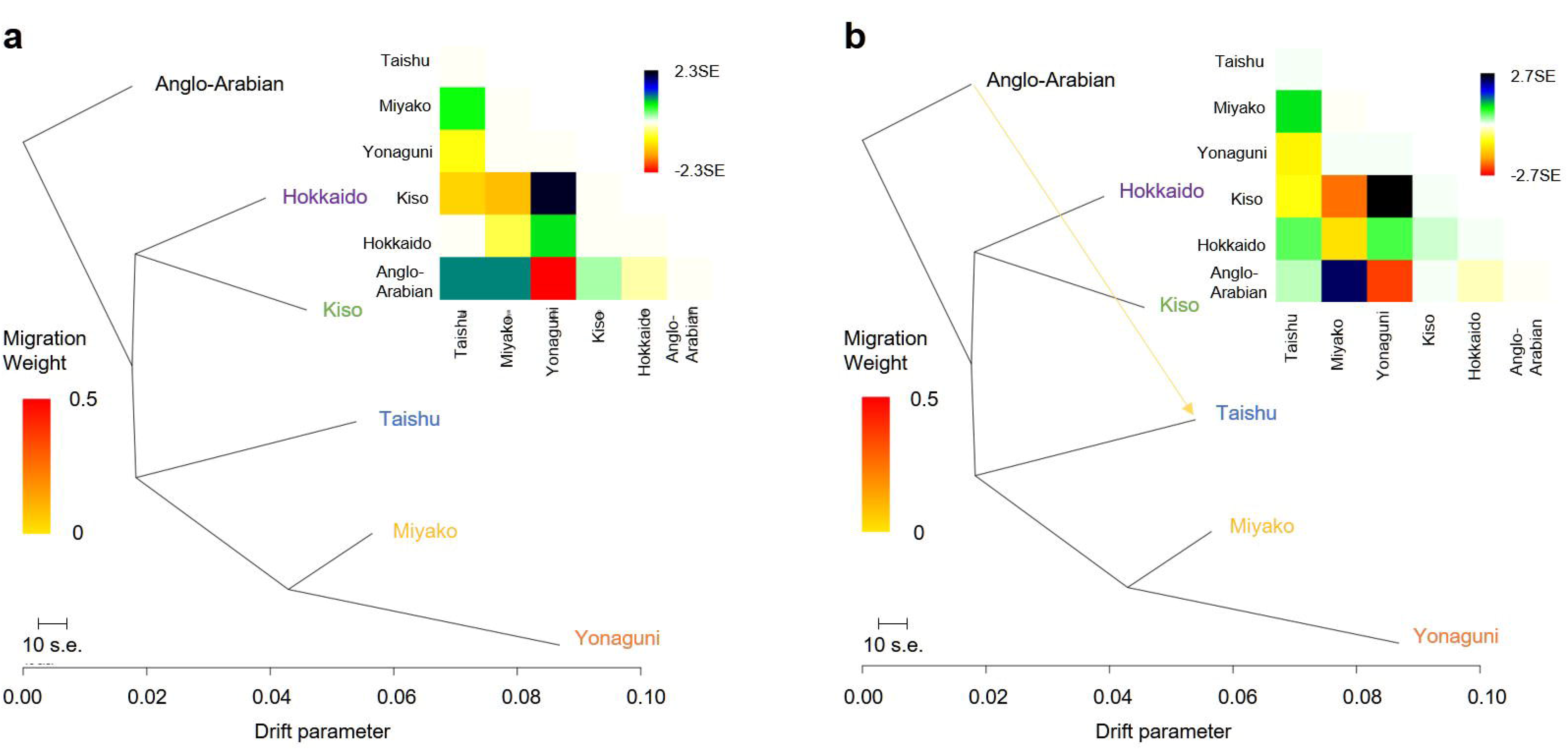
Results of TreeMix analysis of 6 Japanese native horse breeds. Maximum-likelihood trees and the matrices of pairwise residuals for a model allowing (a) 0 migration events and (b) 1 migration event from Anglo-Arabian to Taishu. We estimated that the current Taishu population would have 4% of their ancestry from Anglo-Arabian. Large positive values in the residual matrix indicate a poor fit for the respective pair of populations. Edges representing mixture events are colored according to the weight of the inferred edge.

### Genetic introgression at the individual level - STRUCTURE analysis -

To estimate the degree of genetic introgression from Anglo-Arabian individuals to Taishu at individual level, we used 9609 loci with 1 SNP and 2 alleles in the STRUCTURE analysis. When we used genomic data from all individuals, based on *Ln P*(K) and delta K (EVANNO et al. 2005), we determined that the optimal value of K was 2 (Fig. 3a). One cluster included Taishu, another included Miyako and Yonaguni. Kiso, Hokkaido, and Anglo-Arabian consisted of 1/3 Taishu cluster and 2/3 Miyako-Yonaguni cluster. When K = 5 and 6, results showed an Anglo-Arabian cluster, and that all breeds were grouped individually; however, some Taishu were partially included in the same cluster as Anglo-Arabian. Then, we conducted additional structure analysis using genomic data from only Taishu and Anglo-Arabian. Based on *Ln P*(K) and delta K (EVANNO et al. 2005), we determined that the optimal value of K was 2 (Fig. 3b). When K = 3, results showed an Anglo-Arabian cluster. In both analyses, we observed 10 individuals who comprised 0.1–10 % of the Anglo-Arabian cluster. However, this degree comprising Anglo-Arabian cluster was detected in all 5 Japanese native breeds (Fig. 3a and Fig. 3b).

**Fig. 3.**
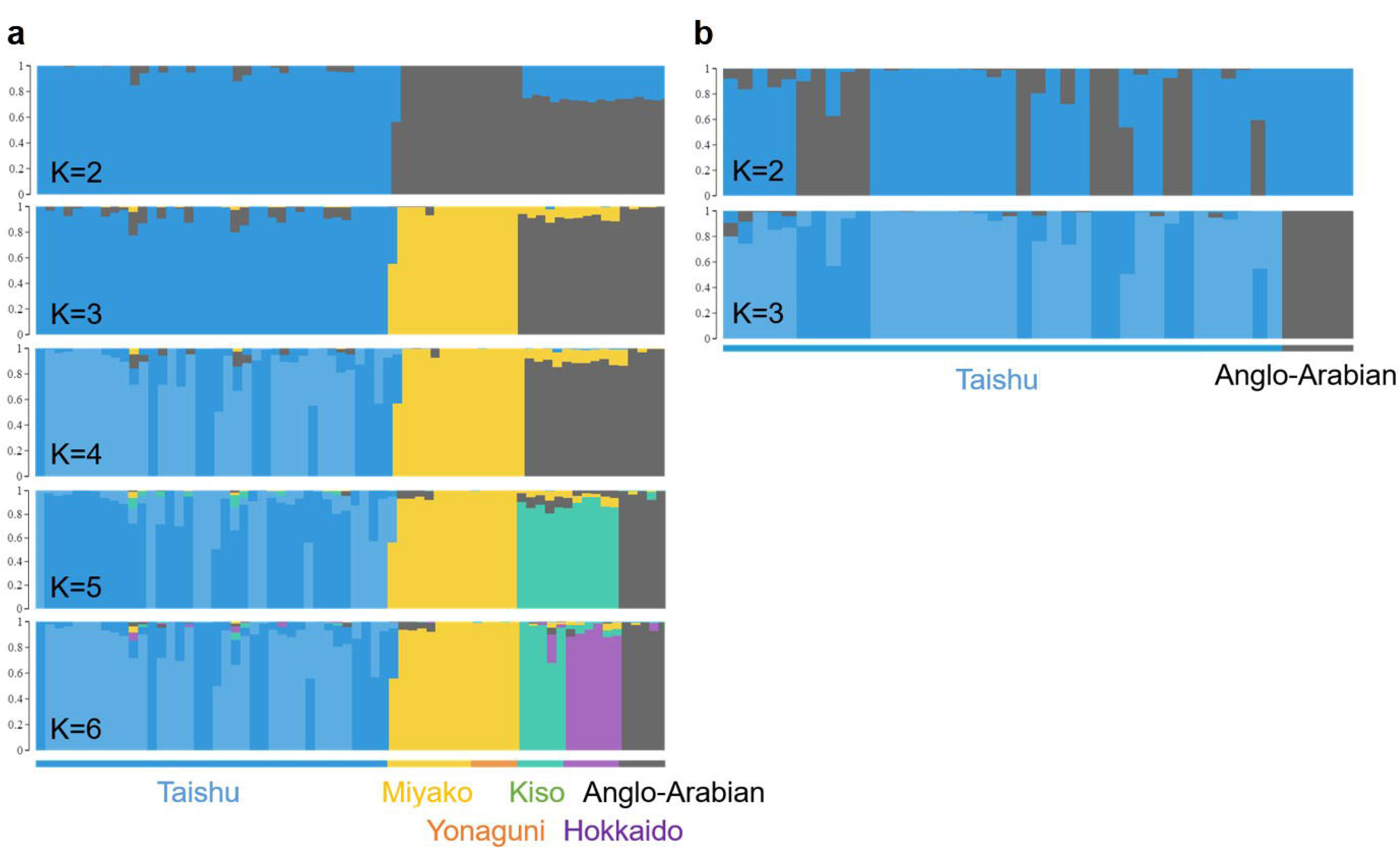
Genetic clustering using STRUCTURE with an admixture model. (a) Structure results for 5 Japanese horse breeds and Anglo-Arabian from K = 2 to 6. (b) Structure results for Taishu and Anglo-Arabian from K = 2 and K = 3.

### Genetic introgression at the individual level - ABBA/BABA tests for each Taishu individuals -

To confirm that the genetic introgression from Anglo-Arabian to Taishu is more often than in other native Japanese breeds, we conducted four ABBA/BABA tests for all Taishu individuals (Fig. 4 and Supplemental Material S6). The ABBA/BABA test calculates the proportion of ABBA and BABA patterns. An excess of any of these patterns indicates the genetic introgression that can be detected using Patterson’s D statistic (Green et al. 2010). If D is significantly different from 0, then the null hypothesis of no genetic introgression is rejected. When using the SNPs of Taishu and 3 of the native Japanese breeds, without Yonaguni as a sister species, almost all individuals had negative Patterson’s D. Thus, the genetic introgression from Anglo-Arabian to Taishu did not occur as often as it did in the other 3 breeds. On the other hand, genetic migration from Anglo-Arabian into Taishu occurred more often than that into Yonaguni. If genetic introgression from Anglo-Arabian into Taishu during WWII has left a signature in Taishu individuals, then Patterson’s D should be positive when using SNPs of both Miyako and Yonaguni as non-introgressed breeds.

**Fig. 4.**
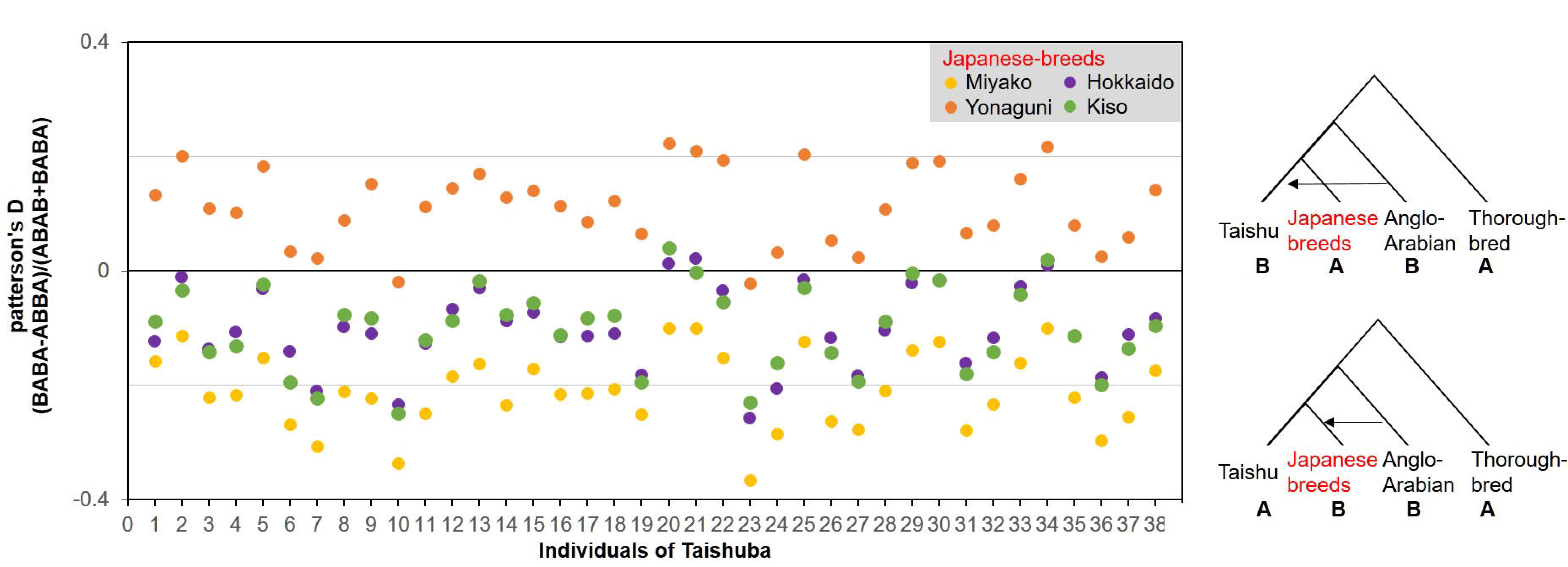
Patterson’s D statistic to test for genetic introgression at the individual level. Positive values indicate gene flow from Anglo-Arabian into Taishu, while negative values indicate gene flow from Anglo-Arabian to other Japanese native breeds. Exact values are shown in Supplemental Material S6.

### Genetic introgression at chromosomal and regional levels - ABBA/BABA tests for each chromosome -

After a genetic introgression event, it is possible that the genetic variations from introgression partially remain in the genome of successive generations of Taishu. Thus, the signature of genetic introgression was not detected using individual data of whole genome SNPs. According to the preceding analysis, it is possible that genetic introgression from Anglo-Arabian into Taishu remained in small regions of Taishu genomes. Therefore, we calculated Patterson’s D for each chromosome per sample to detect the signatures of genetic introgression from Anglo-Arabian to Taishu at each chromosome (Fig. 5 and Supplemental Material S7). The results of the ABBA/BABA tests at the chromosomal level showed negative Patterson’s D for almost all chromosomes when using the SNPs of Taishu and 3 of the native Japanese breeds, without Yonaguni, as a sister species. However, Patterson’s D for chromosomes 21 and 24 were positive in all four patterns of the ABBA/BABA tests (Fig. 5). This strongly indicated that chromosomes 21 and 24 had retained the genetic introgression.

**Fig. 5.**
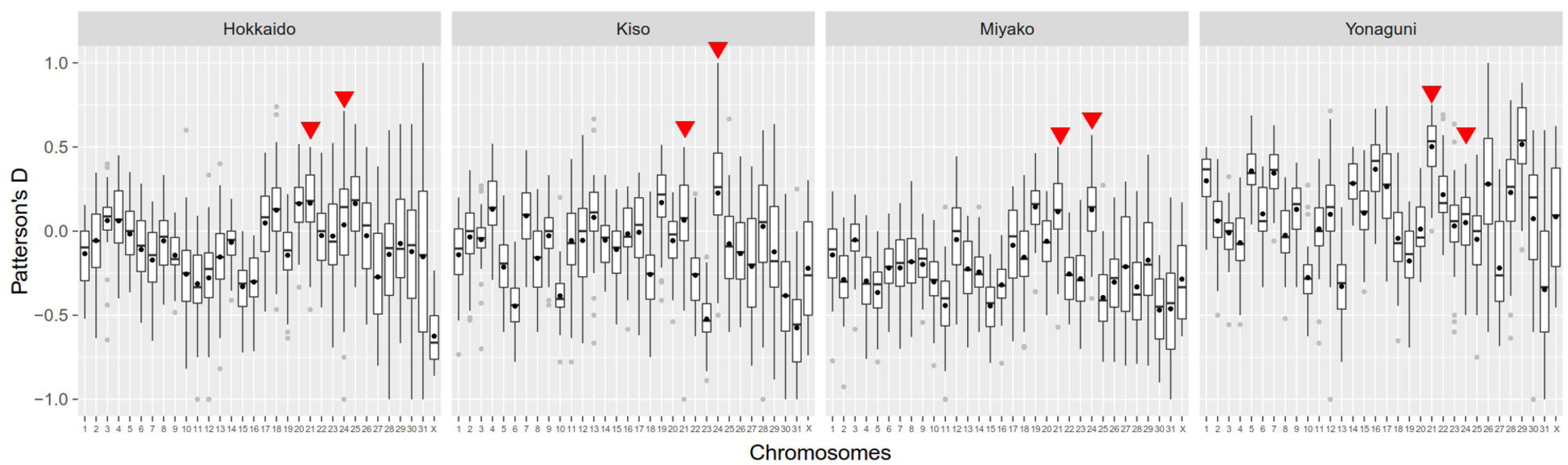
Patterson’s D statistic to test for genetic introgression at the chromosome and region levels. Patterson’s D statistics for each chromosome. Red triangles indicated mean D >0 among four patterns of ABBA/BABA tests. Black circles, black bars, and gray circles indicated mean, median, and outliers of D, respectively.

### Genetic introgression at chromosomal and regional levels - ABBA/BABA tests for regions at Chr 21 and Chr 24 -

It is assumed that because of incomplete genetic recombination after genetic introgression, the signatures of recent genetic introgression (i.e., during WWII) combined on the each of the chromosomes. To confirm that the signatures on the each of Chr 21 and Chr 24 combined with each other, we calculated Patterson’s D for Chr 21 and Chr 24 using 10 non-overlapping SNPs (roughly 10 kb regions) sliding window analysis (Supplemental Material S3, S8 and S9). The regions with positive D were calculated in approximately half of the chromosomes and were on separate chromosomes.

### Genetic introgression at the locus level - Shared SNPs among 5 Japanese native breeds and Anglo-Arabian breed -

We counted shared SNPs between Anglo-Arabian to Taishu as the potential introgressed SNPs. The number of breed-specific SNPs is shown in Fig. 6. In current Taishu population, 8 504 loci showed variations. There were 554 loci with Taishu-specific SNPs, which should be retained in conservation plans for this breed. For SNPs that were shared between Taishu and Anglo-Arabian, there were 961 loci that had “potential” introgressed SNPs, which could have reduced the frequency of the SNPs in Taishu. Another 6 988 loci were considered as ancestral variations, which is a common ancestor of all six breeds in this study and should be retained as much as possible.

**Fig. 6.**
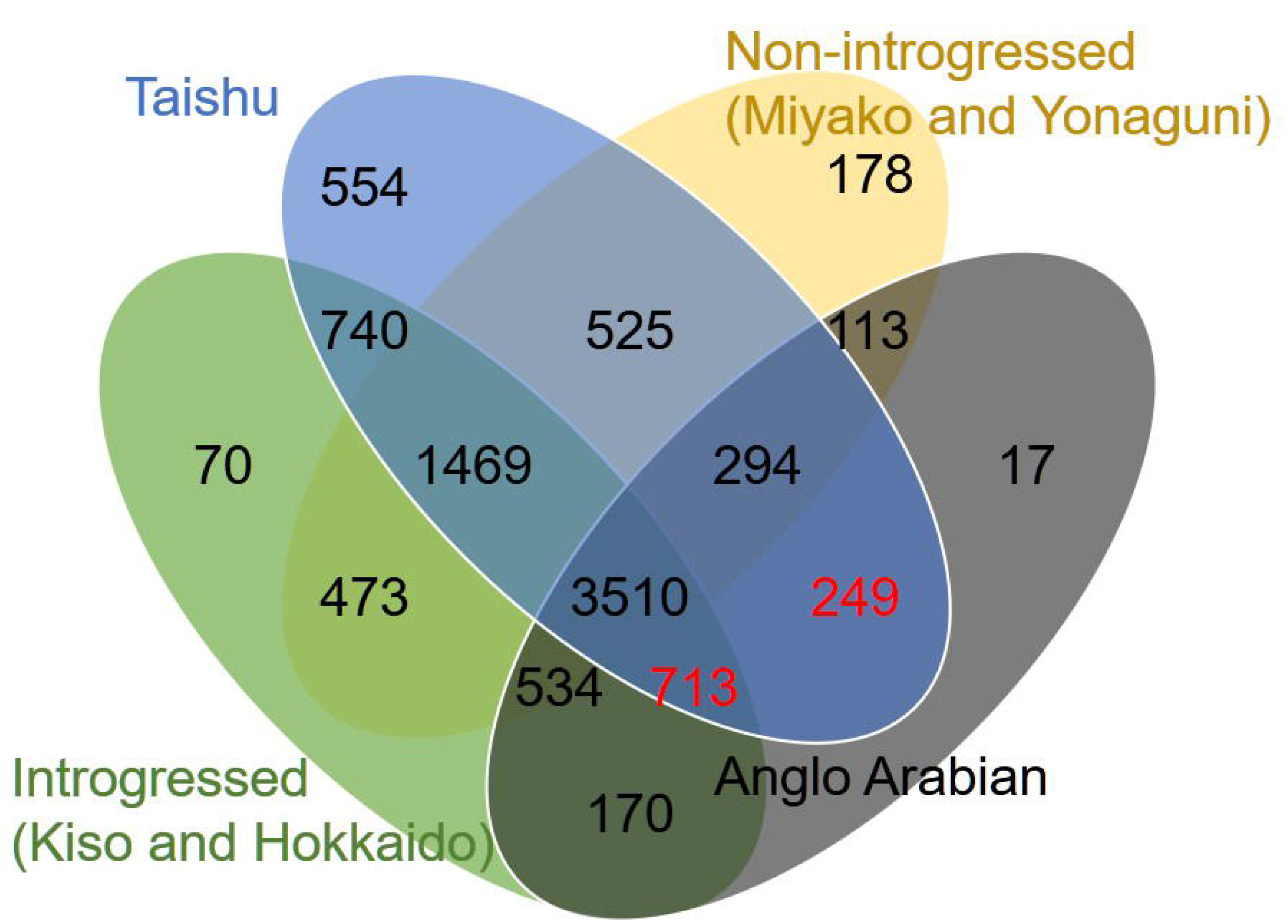
SNP configurations. SNPs of Taishu, non-introgressed breeds (including Miyako and Yonaguni), introgressed breeds (including Kiso and Hokkaido), and Anglo-Arabian are represented by blue, yellow, green, and grey, respectively.

The number of Taishu-specific SNPs was almost correlated with the size of the chromosome (All SNPs = 31.6 + 2.9 x genome size of each chromosome (Mb), R^2^ = 0.88, SE = 38.6, P < 0.001), but the positive correlation between the potential introgressed SNPs and chromosome size was lower than that for all SNPs (introgressed SNPs = 0.37 + 2.5 x genome size of each chromosome (Mb), R^2^ = 0.68, SE = 9.12, *P* < 0.001). For example, there were a low number of potential introgressed SNPs on chromosome X, which is the second longest chromosome in the horse genome (Supplemental Material S4). We counted the number of “potential” introgression SNPs at 7 loci on chromosome 21 (9.9% of all shared SNPs on Chr 21) and 19 loci on chromosome 24 (38.8% of all shared SNPs on Chr 24). Chromosomes 21 and 24 which were estimated as introgressed chromosomes by the preceding analysis did not show a substantial number of shared SNPs more than other chromosomes (Supplemental Material S4).

### Genetic introgression at the locus level - Shared SNPs between Taishu and Anglo-Arabian on high linkage disequilibrium -

Patterson’s D differed depending on the sister breed of the four Japanese native breeds (Fig. 4 and 5); therefore, we conducted another test for recent genetic introgression without using the data from the four Japanese breeds. The genetic introgression from Anglo-Arabian into Taishu occurred relatively recently; therefore, we expected that recent-introgressed SNPs should be in the genomic regions of high linkage disequilibrium (high LD). We counted the shared SNPs in the high LD regions. Of the potential introgressed SNPs, 36 were in high LD regions (10% of all potential introgressed SNPs in >10 kb high LD regions) in 38 Taishu individuals. The frequency of the potential introgressed SNPs in the high LD regions did not significantly affect the frequency of all SNPs in the high LD regions (2-sample test for equality of proportions with continuity correction, χ = 2.03, df = 1, *p* = 0.154).

### Genetic introgression at the locus level - Genetic diversity of Japanese native horses -

We calculated the nucleotide diversity (*π*) of 5 Japanese native horse breeds as the index of genetic diversity. The nucleotide diversity of Taishu was significantly lower than that of the other breeds (Fig. 7, Bonferroni-adjusted Welch’s *t*-tests, p <0.001 each). However, the inbreeding coefficient (*F_IS_*) for Taishu was not lower than that of other breeds (Supplemental Material S5). Taishu had many unique SNPs (Fig. 6), but most of them have low frequency in the current population.

**Fig. 7.**
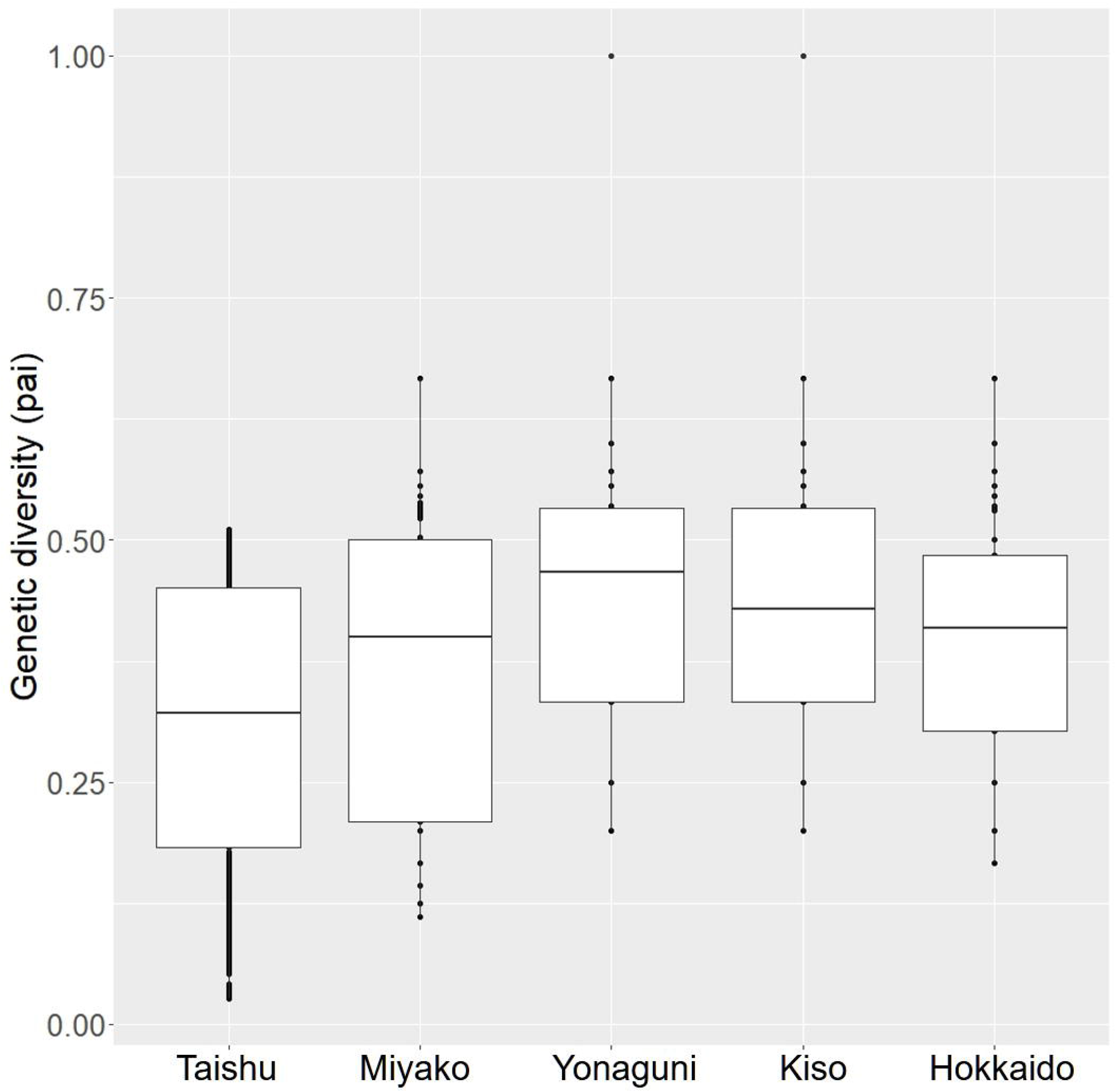
Boxplot of genetic diversity of 5 Japanese native horse breeds. Nucleotide diversity (*π*) of 5 Japanese native horse breeds. Nucleotide diversity of Taishu (n = 38, mean = 0.3075) was significantly lower than that of the other breeds (Miyako: n = 9, mean = 0.3598; Yonaguni: n = 5, mean = 0.4133; Kiso: n = 5, mean = 0.4134; Hokkaido: n = 6, mean = 0.3733).

## Discussion

### Genetic introgression from Anglo-Arabian to Taishu

We conducted the main 8 analyses to confirm genetic introgression from Anglo-Arabian to Taishu and to estimate genetic diversity of Taishu (Table 1). First, we reconstructed the ML phylogeny by using ddRAD-seq markers to confirm admixture between Anglo-Arabian population and Taishu population (Fig. 1 and Table 1). All individuals grouped based on the breeds and this clear pattern of phylogeny corresponded with previous studies (Tozaki et al. 2003). Tozaki et al. (2003) showed that Kiso and Hokkaido were grouped in a sister clade and that Kiso-Hokkaido clade and Taishu were grouped in a sister clade. It is reasonable to group Yonaguni and Miyako in a sister clade because their habitats are geographically very close. Both Yonaguni and Miyako have smaller body sizes compared with other breeds of Japanese native horses. Thus, we considered that the ML phylogeny by using ddRAD-seq markers is reliable. The phylogeny showed that Taishu population has sub-populations, and the genetic distances between Taishu subpopulations were relatively large. One possible cause could be the rapid decrease in population size. Another possible cause could be because of the inconvenience caused by the blockage of movement between the north and south islands, which may have resulted in the differences in the breeding populations on Tsushima island. The genetic distances of both subpopulations of Taishu and Anglo-Arabian were large; therefore, we concluded that this structure was not derived from introgression of the Anglo-Arabian breed with Taishu. Then, we reconstructed using the phylogeny of 16S rRNA. The 16S rRNA phylogeny did not reflect the pattern of breeds (Supplemental Material S2) and this concurred with previous studies that show mtDNA in the horse is highly diverse and does not show the phylogenetic pattern of breeds in European and Asian horses (Achilli et al. 2012). This indicated that using partial DNA sequences was not suitable for detecting introgression because they show biased variation in the genomes. Specific regions of horse genomes have undergone rapid and strong artificial selection while others have maintained the ancestral variations from before domestication (Achilli et al. 2012).

**Table 1.**
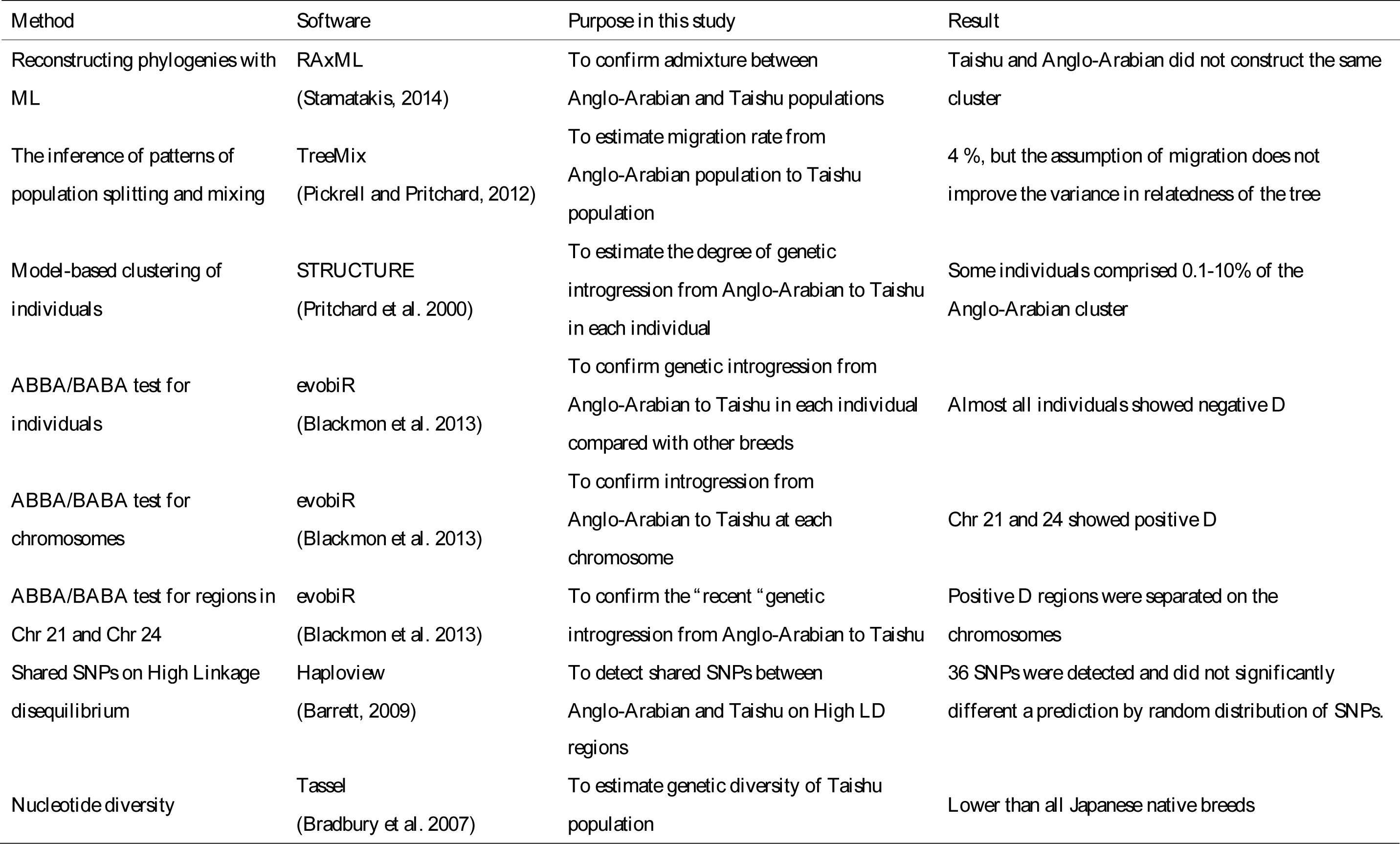
The summary of the analysis and the results in this study.

Second, we estimated the migration rate of Anglo-Arabian population to Taishu population by TreeMix analysis. The results showed that the migration rate was 4 % in the best-explained tree and the assumption that migration from Anglo-Arabian to Taishu did not dramatically improve the variance in relatedness of the tree, indicated that almost no introgression from Anglo-Arabian into Taishu remained in the current population.

Then, we conducted STRUCTURE analysis to assess the degree of genetic introgression at individual level (Fig. 3 and Table 1). The results of STRUCTURE Analysis showed that 10 individuals of all Taishu individuals comprised 0.1–10 % of the Anglo-Arabian cluster. As this degree of Anglo-Arabian clusters were detected in individuals in other Japanese native breeds (Fig. 3a and 3b), it is not enough to conclude that Taishu individuals who comprised the Anglo-Arabian clusters are introgressed individuals.

Subsequently, we conducted another analysis, ABBA/BABA test, to assess the degree of genetic introgression in comparison with 4 other Japanese breeds (Fig. 4 and Table 1). When using the SNPs of Taishu and 3 of the native Japanese breeds, without Yonaguni as a sister species, almost all individuals had negative Patterson’s D, which indicated that the introgression from Anglo-Arabian did not occur in more than 3 breeds. When Yonaguni and Taishu were included, the results indicated that genetic migration from Anglo-Arabian into Taishu occurred more often than into Yonaguni. If recent genetic introgression (i.e., during WWII) from Anglo-Arabian into Taishu has left a signature in Taishu individuals, then Patterson’s D should be positive when using SNPs of non-introgressed breeds (Miyako and Yonaguni). It is difficult to conclude the results of ABBA/BABA test. ABBA/BABA test showed positive D when using Yonaguni as a sister species of Taishu, meaning that genetic introgression from Anglo-Arabian to Taishu was often more than to Yonaguni; however, ABBA/BABA test showed negative D when using Miyako, meaning that genetic introgression from Anglo-Arabian to Taishu was not often more than Miyako, even though both Yonaguni and Miyako were non-introgressed breeds. There are some possible causes of the negative D when using 3 of the native Japanese breeds, without Yonaguni as a sister species. One possibility could be that introgression partially remained in the genome of Taishu individuals and another is that the signatures of introgression was derived from genetic admixture long before genetic introgression during WWII. Therefore, we calculated Patterson’s D for each chromosome per sample (Fig. 5 and Table 1). Patterson’s D for chromosomes 21 and 24 were positive in all four patterns of the ABBA/BABA tests. These results indicated strongly that chromosomes 21 and 24 had retained the genetic introgression. If the signatures of genetic introgression on Chr 21 and Chr 24 were derived from the recent genetic introgression, they must have combined on the each of the chromosomes. To confirm that the signatures on Chr 21 and 24 combined each other, we calculated Patterson’s D at region level. Regions with positive Patterson’s D were on separate chromosomes (Supplemental Material S3). The results suggested that the signatures of “recent” genetic introgression were not found in the current Taishu population.

### “Recent” genetic introgression from Anglo-Arabian to Taishu

We counted the shared SNPs to assess the genetic introgression at loci level and to confirm the signatures of genetic introgression were derived from recent genetic introgression during WWII (Fig. 6). There was a low number of potential introgressed SNPs on chromosome X (Supplemental Material S4). Although mean values of Patterson’s D at chromosome X were negative, chromosomes 21 and 24 that were estimated as introgressed chromosomes by the preceding analysis did not show a substantial amount of shared SNP when compared to other chromosomes (Supplemental Material S4). This suggested that the estimation of genetic introgression using only the number of shared variations is not completely accurate. Then, we counted the shared SNPs in the high LD regions because if the genetic introgression from Anglo-Arabian into Taishu occurred recently, the introgressed SNPs were present in the high LD regions (Table 1). 36 SNPs were in high LD regions (10% of all shared SNPs) in 38 Taishu individuals. The frequency of the shared SNPs in the high LD regions did not significantly affect the frequency of all SNPs in the high LD regions.

These results suggested that although the genetic introgression event between Anglo-Arabian and Taishu was recorded during WWII, the incontestable signatures of the recent genetic introgression on their genome were not detected in the current Taishu population. Therefore, it is likely that the Taishu population has undergone a drastic decrease in size with a hard bottleneck, and the introgressed offspring were not preferred by residents on the Tsushima islands. It is possible that many Anglo-Arabian offspring could not tolerate the environment on the islands. The number of Taishu individuals on Tsushima island was 2405 in 1952 (Hayashida, 1972); thus, the population has declined to about 1/60 of its former size. Moreover, the residents of Tsushima island prefer individuals with a smaller body size that are more suitable for agricultural purposes (Hayashida, 1972). Also, Taishu has been maintained on the island without forage, indicating that Taishu has a higher tolerance to low nutrition (Hayashida, 1972), whereas the offspring of Anglo-Arabian and Taishu might have weaker resistance.

### Genetic diversity of Taishu

The nucleotide diversity of Taishu was significantly lower than that of the other breeds (Fig. 7 and Table 1), despite the genetic introgression from Anglo-Arabian, which increased the nucleotide diversity of this breed. This is consistent with the results of other tests of introgression (Table 1). The inbreeding coefficient (*F_IS_*) for Taishu was not higher than that of other breeds (Supplemental Material S5), indicating that the low nucleotide diversity of Taishu was not due to a failure of recent artificial breeding. There are many unique SNPs in Taishu (Fig. 6), but they are less frequent in the current population and thus, will be lost. In general, low genetic diversity affects the long-term potential for survival of populations (Bouzat, 2010), and individual fitness because of decreased sperm quality (Hedrick and Fredrickson, 2010), reduced litter size (Hedrick and Fredrickson, 2010), increased mortality of juveniles (RALLS et al. 1988), and increased susceptibility to diseases and parasites (Coltman et al. 1999). Although Taishu has undergone genetic introgression from Anglo-Arabian, there were no, or very few, signatures of recent genetic introgression in the current Taishu population. This suggests that the decrease in genetic diversity is a more urgent issue than the removal of genetic introgression.

### The utility of ddRAD-Seq for native livestock

Population genetic approaches using ddRAD-Seq can distinguish the breeds of 5 native Japanese horses that have few genetic differences and can evaluate the genetic introgression status and genetic diversity of breeds. In this study, the phylogeny by using ddRAD-seq markers was consistent with the results from Tozaki et al. (2013), suggesting that ddRAD-seq provides reliable data. ddRAD-seq can provide variable genome wide markers for native livestock that do not have SNP chips or genomic information.

By applying ddRAD-Seq to native livestock, we could have conducted further downstream analysis. Genome-wide markers can provide two methods of analysis: all markers together and markers divided into regions. After several generations, it is possible that the ability to detect introgression is weaker using all genome-wide markers together, because the introgressed regions represent only parts of the genome. In fact, the signatures of introgression in Taishu were found at the chromosomal level and not at the individual level. However, “recent” genetic introgression was not supported by other analyses, and we concluded that the signatures of introgression reflected events older than the introgression event during WWII. Although the signatures of recent genetic introgression events were not detected in many of the analyses, many of the shared SNPs between Taishu and Anglo-Arabian were detected. This indicated that defining genetic introgression using only shared SNPs might lead to overestimation of genetic introgression, while using an insufficient number of, and unevenly distributed, markers also carries a risk of misestimation of introgression. Thus, to detect both the presence and the degree of introgression in native livestock, we need to use both shared SNPs and population genetic approaches using large numbers of genome-wide markers.

## Acknowledgments

We thank the conservation organizations of Taishu, Kiso, Miyako, Yonaguni, and Goryo Bokujyo in the imperial household agency for providing samples, Fumie Kobayashi and Satoko Kondo for their help with the experiments, Naomi Niwa for taking care of paperwork, and Yumie Shinohara for help with all of this research. Funding: This work was supported by the Research Institute for Food and Agriculture of Ryukoku University.

## Reference

Achilli, A., Olivieri, A., Soares, P., Lancioni, H., Kashani, B.H., Perego, U.A., Nergadze, S.G., Carossa, V., Santagostino, M., Capomaccio, S., Felicetti, M., Al-Achkar, W., Penedo, M.C.T., Verini-Supplizi, A., Houshmand, M., Woodward, S.R., Semino, O., Silvestrelli, M., Giulotto, E., Pereira, L., Bandelt, H.-J., Torroni, A., 2012. Mitochondrial genomes from modern horses reveal the major haplogroups that underwent domestication. Proc. Natl. Acad. Sci. USA 109, 2449–2454. doi: 10.1073/pnas.1111637109

Baird, N.A., Etter, P.D., Atwood, T.S., Currey, M.C., Shiver, A.L., Lewis, Z.A., Selker, E.U., Cresko, W.A., Johnson, E.A., 2008. Rapid SNP Discovery and Genetic Mapping Using Sequenced RAD Markers. PLoS ONE 3, e3376–7. doi: 10.1371/journal.pone.0003376

Barrett, J.C., 2009. Haploview: Visualization and Analysis of SNP Genotype Data. Cold Spring Harb. Protoc. 2009, pdb.ip71–pdb.ip71. doi: 10.1101/pdb.ip71

Blackmon, H., Adams, R.H., Blackmon, M.H., 2013. Package “evobiR.”

Bolger, A.M., Lohse, M., Usadel, B., 2014. Trimmomatic: a flexible trimmer for Illumina sequence data. Bioinformatics 30, 1–7. doi: 10.1093/bioinformatics/btu170

Bouzat, J.L., 2010. Conservation genetics of population bottlenecks: the role of chance, selection, and history. Conserv. Genet. 11, 463–478. doi: 10.1007/s10592-010-0049-0

Bradbury, P.J., Zhang, Z., Kroon, D.E., Casstevens, T.M., Ramdoss, Y., Buckler, E.S., 2007. TASSEL: Software for association mapping of complex traits in diverse samples. Bioinformatics 23:2633-2635.

Browning, B.L., Browning, S.R., 2016. Genotype Imputation with Millions of Reference Samples. Am. J. Hum. Genet. 98, 116–126. doi: 10.1016/j.ajhg.2015.11.020

Catchen, J., Hohenlohe, P.A., Bassham, S., Amores, A., Cresko, W.A., 2013. Stacks: an analysis tool set for population genomics. Mol. Ecol. 22, 3124–3140. doi: 10.1111/mec.12354

Chattopadhyay, B., Garg, K.M., Kumar, A.K.V., Doss, D.P.S., Rheindt, F.E., Kandula, S., Ramakrishnan, U., 2016. Genome-wide data reveal cryptic diversity and genetic introgression in an Oriental cynopterine fruit bat radiation. BMC Evol. Biol. 16, 41. doi: 10.1186/s12862-016-0599-y

Coltman, D.W., Pilkington, J.G., Smith, J.A., Pemberton, J.M., 1999. Parasitec against inbred soay sheep in a free living island population. Evolution 53, 1259–1267. doi: 10.1111/j.1558-5646.1999.tb04538.x

Combosch, D.J., Vollmer, S.V., 2015. Trans-Pacific RAD-Seq population genomics confirms introgressive hybridization in Eastern Pacific Pocillopora corals. Mol. Phylogenet. Evol. 88, 154–162. doi: 10.1016/j.ympev.2015.03.022

Earl, D.A., vonHoldt, B.M., 2012. STRUCTURE HARVESTER: a website and program for visualizing STRUCTURE output and implementing the Evanno method. Conserv. Genet. Resour. 4, 359–361. doi: 10.1007/s12686-011-9548-7

Evanno, G., Regnaut, S., Goudet, J., 2005. Detecting the number of clusters of individuals using the software structure: a simulation study. Mol. Ecol. 14, 2611–2620. doi: 10.1111/j.1365-294X.2005.02553.x

Fitzpatrick, B.M., Johnson, J.R., Kump, D.K., Smith, J.J., Voss, S.R., Shaffer, H.B., 2010. Rapid spread of invasive genes into a threatened native species. Proc. Natl. Acad. Sci. USA 107, 3606–3610. doi: 10.1073/pnas.0911802107

Futschik, A., Schlotterer, C., 2010. The Next Generation of Molecular Markers From Massively Parallel Sequencing of Pooled DNA Samples. Genetics 186, 207–218. doi: 10.1534/genetics.110.114397

Gabriel, S.B., Schaffner, S.F., Nguyen, H., Moore, J.M., Roy, J., Blumenstiel, B., Higgins, J., DeFelice, M., Lochner, A., Faggart, M., Liu-Cordero, S.N., Rotimi, C., Adeyemo, A., Cooper, R., Ward, R., Lander, E.S., Daly, M.J., Altshuler, D., 2002. The Structure of Haplotype Blocks in the Human Genome. Science 296, 2225–2229. doi: 10.1126/science.1069424

Hanotte, O., Dessie, T., Kemp, S., 2010. Time to Tap Africa’s Livestock Genomes. Science 1640–1641. doi: 10.1126/science.1186254

Hayashida, S., 1972. Native horse in Tsushima island “Taishuba.” Japan Racing Association, Minato-ku.

Hedrick, P.W., Fredrickson, R., 2010. Genetic rescue guidelines with examples from Mexican wolves and Florida panthers. Conserv. Genet. 11, 615–626. doi: 10.1007/s10592-009-9999-5

Larkin, M.A., Blackshields, G., Brown, N.P., Chenna, R., McGettigan, P.A., McWilliam, H., Valentin, F., Wallace, I.M., Wilm, A., Lopez, R., Thompson, J.D., Gibson, T.J., Higgins, D.G., 2007. Clustal W and Clustal X version 2.0. Bioinformatics 23, 2947–2948. doi: 10.1093/bioinformatics/btm404

Mardis, E.R., 2008. Next-Generation DNA Sequencing Methods. Annu. Rev. Genom. Human Genet. 9, 387–402. doi: 10.1146/annurev.genom.9.081307.164359

McCue, M.E., Bannasch, D.L., Petersen, J.L., Gurr, J., Bailey, E., Binns, M.M., Distl, O., Guérin, G., Hasegawa, T., Hill, E.W., Leeb, T., Lindgren, G., Penedo, M.C.T., Røed, K.H., Ryder, O.A., Swinburne, J.E., Tozaki, T., Valberg, S.J., Vaudin, M., Lindblad-Toh, K., Wade, C.M., Mickelson, J.R., 2012. A High Density SNP Array for the Domestic Horse and Extant Perissodactyla: Utility for Association Mapping, Genetic Diversity, and Phylogeny Studies. PLoS Genet. 8, e1002451–14. doi: 10.1371/journal.pgen.1002451

Nielsen, R., Paul, J.S., Albrechtsen, A., Song, Y.S., 2011. Genotype and SNP calling from next-generation sequencing data. Nat. Rev. Genet. 12, 443–451. doi: 10.1038/nrg2986

Peterson, B.K., Weber, J.N., Kay, E.H., Fisher, H.S., Hoekstra, H.E., 2012. Double Digest RADseq: An Inexpensive Method for De Novo SNP Discovery and Genotyping in Model and Non-Model Species. PLoS ONE 7, e37135–11. doi: 10.1371/journal.pone.0037135

Pickrell, J.K., Pritchard, J.K., 2012. Inference of Population Splits and Mixtures from Genome-Wide Allele Frequency Data. PLoS Genet. 8, e1002967. doi: 10.1371/journal.pgen.1002967

Poland, J.A., Rife, T.W., 2012. Genotyping-by-Sequencing for Plant Breeding and Genetics. Plant Genome 5, 92–11. doi: 10.3835/plantgenome2012.05.0005

Pritchard, J.K., Stephens, M., Donnelly, P., 2000. Inference of Population Structure Using Multilocus Genotype Data. Genetics 155, 945–959. doi: 10.1016/0379-0738(94)90222-4

Ralls, K., Ballou, J.D., Templeton, A., 1988. Estimates of Lethal Equivalents and the Cost of Inbreeding in Mammals. Conserv.Biol. 2, 185–193. doi: 10.1111/j.1523-1739.1988.tb00169.x

Ramasamy, R.K., Ramasamy, S., Bindroo, B.B., Naik, V.G., 2014. STRUCTURE PLOT: a program for drawing elegant STRUCTURE bar plots in user friendly interface. SpringerPlus 3, 431.

Green, R.E., Krause, J., Briggs, A.W., Maricic, T., Stenzel, U., Kircher, M., Patterson, N., Li, H., Zhai, W., Fritz, M.H.Y., Hansen, N.F., 2010. A Draft Sequence of the Neandertal Genome. Science 328, 710–722.

Rischkowsky, B., Pilling, D., 2007. The State of the World’s Animal Genetic Resources for Food and Agriculture. Food & Agriculture Org.; 2007.

Ryan, M.E., Johnson, J.R., Fitzpatrick, B.M., 2009. Invasive hybrid tiger salamander genotypes impact native amphibians. Proc. Natl. Acad. Sci. USA 106, 11166–11171. doi:10.1073/pnas.0902252106

Schaefer, R.J., Schubert, M., Bailey, E., Bannasch, D.L., Barrey, E., Bar-Gal, G.K., Brem, G., Brooks, S.A., Distl, O., Fries, R., Finno, C.J., Gerber, V., Haase, B., Jagannathan, V., Kalbfleisch, T., Leeb, T., Lindgren, G., Lopes, M.S., Mach, N., da Câmara Machado, A., MacLeod, J.N., McCoy, A., Metzger, J., Penedo, C., Polani, S., Rieder, S., Tammen, I., Tetens, J., Thaller, G., Verini-Supplizi, A., Wade, C.M., Wallner, B., Orlando, L., Mickelson, J.R., McCue, M.E., 2017. Developing a 670k genotyping array to tag ~2M SNPs across 24 horse breeds. BMC Genomics 18, 565. doi:10.1186/s12864-017-3943-8

Scherf, B.D., Pilling, D., 2015. The Second Report on the State of the World’s Animal Genetic Resources for Food and Agriculture.

Stamatakis, A., 2014. RAxML version 8: a tool for phylogenetic analysis and post-analysis of large phylogenies. Bioinformatics 30, 1312–1313. doi:10.1093/bioinformatics/btu033

Tozaki, T., Takezaki, N., Hasegawa, T., Ishida, N., Kurosawa, M., Tomita, M., Saitou, N., Mukoyama, H., 2003. Microsatellite Variation in Japanese and Asian Horses and Their Phylogenetic Relationship Using a European Horse Outgroup. J. Hered. 94, 374–380. doi:10.1093/jhered/esg079

Yasugi, M., Tezuka, A., Nagano, A.J., 2018. Stacksbinder: online tool for visualizing and summarizing Stacks output to aid filtering of SNPs identified using RAD sequencing. Conserv Genet Resour https:––doi.org–10.1007–s12686–018–1050–z

